# Scalable and Accurate Drug–target Prediction Based on Heterogeneous Bio-linked Network Mining

**DOI:** 10.1101/539643

**Authors:** Nansu Zong, Rachael Sze Nga Wong, Victoria Ngo, Yue Yu, Ning Li

**Affiliations:** Department of Health Sciences Research, Mayo Clinic, 205 3^rd^ Ave SW, Rochester, MN, 55905; Department of Bioengineering, UC, San Diego, 9500 Gilman Drive, San Diego, CA 92093-0412; Betty Irene Moore School of Nursing, UC, Davis, 2570 48^th^ Street, Sacramento, CA, 95817; Scripps Research Institute, 10550 North Torrey Pines Road, San Diego, CA, 92037

## Abstract

**Motivation:** Despite the existing classification- and inference-based machine learning methods that show promising results in drug-target prediction, these methods possess inevitable limitations, where: 1) results are often biased as it lacks negative samples in the classification-based methods, and 2) novel drug-target associations with new (or isolated) drugs/targets cannot be explored by inference-based methods. As big data continues to boom, there is a need to study a scalable, robust, and accurate solution that can process large heterogeneous datasets and yield valuable predictions.

**Results:** We introduce a drug-target prediction method that improved our previously proposed method from the three aspects: 1) we constructed a heterogeneous network which incorporates 12 repositories and includes 7 types of biomedical entities (#20,119 entities, # 194,296 associations), 2) we enhanced the feature learning method with Node2Vec, a scalable state-of-art feature learning method, 3) we integrate the originally proposed inference-based model with a classification model, which is further fine-tuned by a negative sample selection algorithm. The proposed method shows a better result for drug–target association prediction: 95.3% AUC ROC score compared to the existing methods in the 10-fold cross-validation tests. We studied the biased learning/testing in the network-based pairwise prediction, and conclude a best training strategy. Finally, we conducted a disease specific prediction task based on 20 diseases. New drug-target associations were successfully predicted with AUC ROC in average, 97.2% (validated based on the DrugBank 5.1.0). The experiments showed the reliability of the proposed method in predicting novel drug-target associations for the disease treatment.

## 1 Introduction

Drugs are substances designed to be utilized in the diagnosis, treatment, mitigation or prevention of diseases and its associated symptoms. They can produce physiological effects by binding to a “target” or multiple “targets” to enhance or inhibit functions carried out by these “targets” (Olayan, et al., 2017; Santos, et al., 2017). Exploring novel drug-target interactions (DTI) plays a crucial role in drug development, which facilitate the studying of drug action, disease pathology and drug side effects (Liu, et al., 2015; Santos, et al., 2017; Yuan, et al., 2016). Despite the availability of a variety of biological assays, experimental prediction remains laborious and expensive (Liu, et al., 2015; Wang, et al., 2013), which drives the biochemical experimentation (in vitro) to focus on some particular families of ‘druggable’ proteins, while the potentially larger number of small molecules are rarely systematically screened for paring (Vogt and Mestres, 2010; Yıldırım, et al., 2007). Consequently, in order to lower the overall costs and uncover more potential screening targets, computational (in silico) methods have become popular and are commonly applied for poly-pharmacology and the drugs repurposing in drug development (Cheng, et al., 2012; Ding, et al., 2013).

Early computational methods attempted to predict associations by docking simulations based on the 3D structure of target proteins, or by comparing a query ligand with the ligands of target proteins. These methods were hindered by the lack of ample data, such as 3D structures of proteins or ligands with target proteins (Liu, et al., 2015; Olayan, et al., 2017; Yuan, et al., 2016). To provide the more robust solutions (Cheng, et al., 2007; Zhu, et al., 2005), machine learning algorithms were applied to predict drug–target associations. These methods can generally be categorized into two types: 1) classification-based methods, and 2) inference-based methods (Ding, et al., 2013; Jacob and Vert, 2008; Klipp, et al., 2010). The features used as the input for the diverse classification models play an important role, and the features are generated from three types, 1) chemical structures/fingerprint of drugs and genomic sequence for proteins (Feng, et al., 2018; Liu, et al., 2015; Peng, et al., 2017; Wen, et al., 2017; Yuan, et al., 2016), 2) neighborhood or structure information for vertices (e.g., drugs or targets) in a heterogeneous network structure (Emig, et al., 2013), 3) similarity matrix. It is important to note that similarities are not feature vectors (Ding, et al., 2013). The similarities are directly used as input in the classifications methods, such as kernel regression (Yamanishi, et al., 2008), Bipartite Local Method (BLM) (Bleakley and Yamanishi, 2009), Pairwise Kernel Method (PKM) (Jacob and Vert, 2008), Laplacian regularized least squares (LapRLS) (Xia, et al., 2009). Hence these methods are categorized into feature-base methods. Additionally, these features can be combined to achieve a better result (Olayan, et al., 2017; Yuan, et al., 2016). Different from the binary judgement made based on the classification models, inference-based models often utilize interactions, similarities and correlations between drugs and targets to predict a confidence score for a potential drug-target association. In general, the methods can be categorized to: 1) “guilt-by-association”-based (Alaimo, et al., 2013; Cheng, et al., 2012; Wang, et al., 2013; Zong, et al., 2017), 2) random walk-based (Cheng, et al., 2012; Seal, et al., 2015), 3) similarity-based (Cheng, et al., 2012), and 4) statistical analysis-based (Cheng, et al., 2016). Since the information used in the inference-based methods can be easily pulled from networks, heterogeneous networks often serve as the input (Alaimo, et al., 2013; Cheng, et al., 2012; Cheng, et al., 2016; Luo, et al., 2017; Seal, et al., 2015; Wang, et al., 2013; Zong, et al., 2017). Even though the above methods can achieve satisfactory results in drug-target prediction, there are remarkable limitations. For classification methods, the unknown drug-target pairs are considered as negative samples and the random selection will inevitably include false negative samples for training, which will bias the prediction results (Chen, et al., 2015; Ding, et al., 2013; Peng, et al., 2017). For the inference-based methods, new drugs/targets that previously do not interact with targets/drugs (a.k.a., isolated nodes) cannot be predicted (Chen, et al., 2015; Cheng, et al., 2012; Seal, et al., 2015). The booming of big data also brings opportunities as well as challenges in drug-target prediction. Therefore, scalable and robust solutions that can process large heterogeneous datasets to yield precious prediction for new drugs/targets need to be studied.

The existing machine learning-based works, including our previously proposed network-based (Linked Tripartite Network) solution (Zong, et al., 2017), have three essentials: 1) heterogeneous data, 2) feature learning methods, and 3) prediction generation methods. Therefore, the improvement for the three essentials are the main focus of our study to advance the prediction. In this paper, we introduce a scalable, robust, and accurate drug-target prediction method that improves our previous proposed method from three aspects: 1) to enable more information to support drug-target prediction, we constructed a comprehensive heterogeneous network which incorporates 12 repositories^1^ and includes 7 types of biomedical entities^2^ (#20,119 entities, # 194,296 associations) as the input data, 2) to improve the prediction with a scalable solution, we enhanced the feature learning method with a state-of-art feature learning method, Node2Vec (Grover and Leskovec, 2016), 3) to improve the robustness by enabling prediction based on new drug/targets, we combined the originally proposed inference-based model (Zong, et al., 2017) with a classification model that is fine-tuned with a negative sample selection algorithm for drug-target association learning. We conducted the 10-fold cross-validation tests and showed that the proposed method achieved a better result for drug–target association prediction: 95.3% AUC ROC score compared to the existing methods. In addition, we studied the biased learning/testing in the network-based pairwise prediction (Chen, et al., 2015; Park and Marcotte, 2012), and suggested the best learning strategy for practice. Finally, to evaluate the prediction for disease treatment, we conducted a disease specific prediction task based on 20 types of diseases3. The results were validated by searching DrugBank 5.1.0 (Wishart, et al., 2017) at June 20th, 2018. The experiments showed the reliability of the proposed method (AUC in average: 97.2%) in predicting novel 75 drug-target associations for the disease treatment.

## 2 Methods

### 2.1 Framework of drug-target prediction based on heterogeneous network

We adapted the network-based drug-target prediction framework our previously proposed (Zong, et al., 2017), which contains three steps (see Supplementary figure 1): 1) data preparation & benchmarking, 2) vertex vectorization, and 3) model training and association prediction. First, a multipartite network that contains the topological interactions of the existing drugs and targets as well as the biological interaction with the related entities is constructed. Then, the vector representation of the drugs and targets are learned based on the topology of the network. Finally, new drug-target associations are predicted and evaluated based on a hybrid model that consists of an inference-based model and a classification model based on the existent drug-target associations and the vector representations of the network vertices learned in step 2.

### 2.2 Data preparation and benchmarking

This study utilized information of the various biological entities that related to drugs and targets to form a multipartite network called Linked Multipartite Network (LMN). LMN includes seven biological entities as the vertices and fourteen associations as the edges, and consists of 20,119 vertices and 194,296 edges in total (Table 1).

**Table 1.** Statistics for the Linked Multipartite Network (LMN)

| Entity | Statistics | Association | Statistics |
| --- | --- | --- | --- |
| Drug | 6,990 | Drug-target | 20,583 |
| Target | 4,267 | Disease-target | 13,205 |
| Disease | 3,260 | Drug-disease | 7,232 |
| Sider effect | 1,695 | Drug-drug | 38,140 |
| Pathway | 379 | Disease-disease | 100 |
| Haplotype | 587 | Target-target | 388 |
| Variant location | 2,941 | Disease-haplotype | 3,632 |
|  |  | Drug-haplotype | 3,112 |
|  |  | Disease-variant location | 12,277 |
|  |  | Drug-variant location | 6,868 |
|  |  | Disease-pathway | 5,057 |

<sup>1</sup> DrugBank, Human Disease Network, Sider, KEGG, PharmGKB, Pubchem, OMIM, Uniprot, HGNC, UMLS, MESH, Bio2rdf
<sup>2</sup> Drug, Target, Disease, Sider effect, Pathway, Haplotype, Variant location
<sup>3</sup> 1) Adenocarcinoma, 2) Anemia, 3) Apert syndrome, 4) Autism, 5) Cirrhosis, 6) Galactosemia, 7) Insomnia, 8) Lung Cancer, 9) Malaria, 10) Obesity, 11) Obsessive Compulsive Disorder, 12) Spinal Muscular Atrophy, 13) Stroke, 14) Bipolar Disorder,

|  |  |  |  |
| --- | --- | --- | --- |
|  |  | Drug-pathway | 3,889 |
|  |  | Target-pathway | 10,349 |
|  |  | Drug-side effect | 69,464 |
| In total | 20,119 | In total | 194,296 |

The backbone elements, 6,990 drugs, 4,267 targets, and 20,583 drug-target associations, are from DrugBank and obtained from the linked data repository Bio2rdf (http://download.bio2rdf.org/#/release/4/). Four biological databases, human disease network, Sider, KEGG, and PharmGKB are used to flesh the network along with other six resources, Pubchem, Uniprot, HGNC, OMIM, UMLS and MESH, used as the reference. The construction of the network in detail please see Supplementary File 1.

### 2.3 Association prediction based on graph embedding

We define a network *G*(*V*, *E*) that includes a set of vertices *V* and a set of edges *E* as our input data, where *V* are multiple types of vertices that at least cover two types, set *D* for drugs and *T* for targets, and *E* are multiple types of edges that connect the vertices. If there is an edge connecting a drug and a target, there is a drug-target association between the two concepts in the existing knowledge. While the absence of an edge between a drug and a target represents the unknown association in our knowledge. Given a such network *G*, our goal is to predict the novel associations (unlabeled) between the drugs in *D* and targets in *T* of *V* in *G*. We modeled the problem of drug-target prediction as, given a network *G* as the training data, classifies an unlabeled drug-target association to two categories, “positive” for existent and “negative” for nonexistent association.

#### 2.3.1 Vectorization

In order to train a model with the input network, we used the graph embedding method to learn the features of the nodes/associations. Node2vec (Grover and Leskovec, 2016) is the state-of-art graph embedding method that vectorizes the vertices of a network based on topology of the network with maximum likelihood optimization. Given a network *G* = (*V*, *E*), Node2vec maximizes the probability of observing the neighborhood *N*(*u*) of each node *u* in the network based on two standard assumptions: 1) the likelihood of neighborhood observation is independent, and 2) two nodes have a symmetric effect over each other for the conditional likelihood of source-neighborhood pair. The detail of Node2vec please see the Supplementary File 2.

#### 2.3.2 VectorizationClassification-based prediction

To represent a pair of drug *d* and target *t* for classification, the vectorized representation for the pair f(*d*, *t*) is formalized based on one of the binary operators, Average, Hadamard, Weighted-L1, and Weighted-L2 (Grover and Leskovec, 2016; Rossi, et al., 2018; Zhou, et al., 2018), on the *i*th vector *f*_*i*_ (*d*, *t*) as,

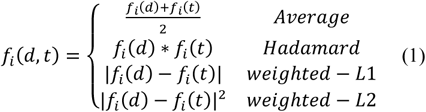

Conventionally, the negative samples (i.e., “false”) used in the classifier training are randomly selected from the huge number of unknown associations (Liu, et al., 2015; Peng, et al., 2017). Therefore, false positives in prediction can be predicted as the unknown positives but labeled as negative (i.e., false negatives) are used in training. We adapted a negative learning method, Spy, to select negative samples from the random selected unknown associations in a certain confidence (Liu, et al., 2002; Peng, et al., 2017).

Given the edges *E* from the network *G*, a set of drug-target associations extracted are considered as positive samples *P*, and a set of randomly selected unknown drug-target associations are *U*. A *s*% positive samples *S* functioned as the spy are randomly chosen to be removed from *P* and join *U* to form *M*. With *M* and *P* as the input, a classifier, I-EM, to classier *M* is obtained by training a Bayesian classifier to maximize the posterior probability of each document in *P*. with I-EM, all the samples that receive a classification score lower than *t* will be considered as the negative, where *t* is determined by how the spy set *S* is classified with I-EM (Liu, et al., 2002). In practice, we take the value of z-score (*P* = 0.01) of score distribution of *S* classification as *t*.

#### 2.3.3 Inferencing-based prediction

We used hybrid-based similarity inference (HBSI) that linearly combines the results of the two popular rule-based inference methods (Cheng, et al., 2012; Yamanishi, et al., 2008), drug-based similarity inference (DBSI) and target-based similarity inference (TBSI). The result of HBSI is calculated as,

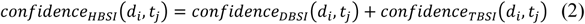

DBSI predicts a drug-target association *s*(*d*_*i*_, *t*_*j*_) if a drug *d*_*i*_ is similar with a drug that has an existing association with a target *t*_*j*_. For a pair of (*d*_*i*_, *t*_*j*_), a confidence score of the pair is calculated as,

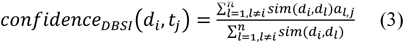

where *sim*(*d*_*i*_, *d*_*l*_) is the similarity between *d*_*i*_ and *d*_*l*_, which is calculated based on vectorizations of *d*_*i*_ and *d*_*l*_. *a*_*l*,*j*_ = 1 if there is an existing association between *d*_*l*_ and *t*_*j*_ otherwise *a*_*l,j*_ = 0 .

Similarly, TBSI is calculated as,

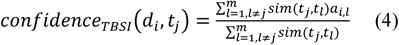

where *sim*(*t*_*j*_, *t*_*l*_) is the similarity between *t*_*j*_ and *t*_*l*_, and *a*_*i,l*_ = 1 if there is an existing association between *d*_*i*_ and *t*_*l*_ otherwise *a*_*i,l*_ = 0.

### 2.4 Methods in comparison

In addition to Node2vec, we also used DeepWalk for the vectorization, which is proved to be the best vectorization method in the inference-based model (Zong, et al., 2017). The two prediction models are used to associate the vectorization model, inferencing-based and classification-based model.

For inferencing-based model, three methods, DBSI, TBSI and HBSI are used to predict an association respectively.

For classification-based model, we adopted four classification methods (Grover and Leskovec, 2016; Liu, et al., 2015; Olayan, et al., 2017; Peng, et al., 2017), SVM, J48, Random Forest, Logistic Regression. Each drug-target pair is formalized with four binary operators, Average, Hadamard, Weighted-L1, and Weighted-L2. In addition, we adopted a semi-classification method, YASTI, which uses the unlabeled test data in training (Driessens, et al., 2006). We associated YASTI with the four classification methods for testing.

In practice, the logistic regression classifier is obtained from the LIBLINEAR library (https://www.csie.ntu.edu.tw/~cjlin/liblinear/), J48 and random forest are obtained from Weka library (https://www.cs.waikato.ac.nz/ml/weka/), SVM is obtained from LIBSVM (https://www.csie.ntu.edu.tw/~cjlin/libsvm/), and YASTI is obtained from collective library (https://github.com/fracpete/collective-classification-weka-package).

### 2.5 Validation and evaluation metrics

We used the convention 10-fold cross-validation for the evaluation (Wang, et al., 2013). To fairly compare the proposed method to the existing ones, we limited the predictions to the drug-target associations containing at least one un-isolated node. To achieve this, we first randomly extracted a set of drug-target associations *A*_*r*_, making sure that no isolated vertices of drugs and targets are created. The remaining associations *A*_*c*_ in the network will be used for training. The associations *A*_*r*_ were randomly partitioned into 10 subsets {*A*1, …., *A*10}. In each test of ten, a subset *A*_*i*_ was used as a gold standard for testing while the nine remaining subsets of *A*_*r*_ as well as *A*_*c*_ were used as the training set.

We used Area Under the Receiver Operating Characteristic Curve (AUC ROC) (Cheng, et al., 2012; Seal, et al., 2015), to assess the quality of the predictions. In practice, AUC ROC scores are calculated by the ROC JAVA library (https://github.com/kboyd/Roc) and Weka evaluation package (Holmes, et al., 1994).

## 3 Results

### 3.1 Comparison with topology-based methods

To comprehensively evaluate the proposed method, we compared the proposed method to the existing network topological-based methods that utilized different solutions in vectorization and association prediction components.

Figure 1 shows that the best result is achieved by the proposed method based on Node2vec (AUC ROC: 95.3%). The proposed method outperformed the two state-of-art solutions, TBSI+Deepwalk: 91.7% and RF+Hadamard+Node2vec: 93.0%, and two proposed variations of the two methods in this study, HBSI+Deepwalk: 91. 9% and YASTI+SVM+Hadamard+Node2vec: 93.1%.

**Figure 1.**
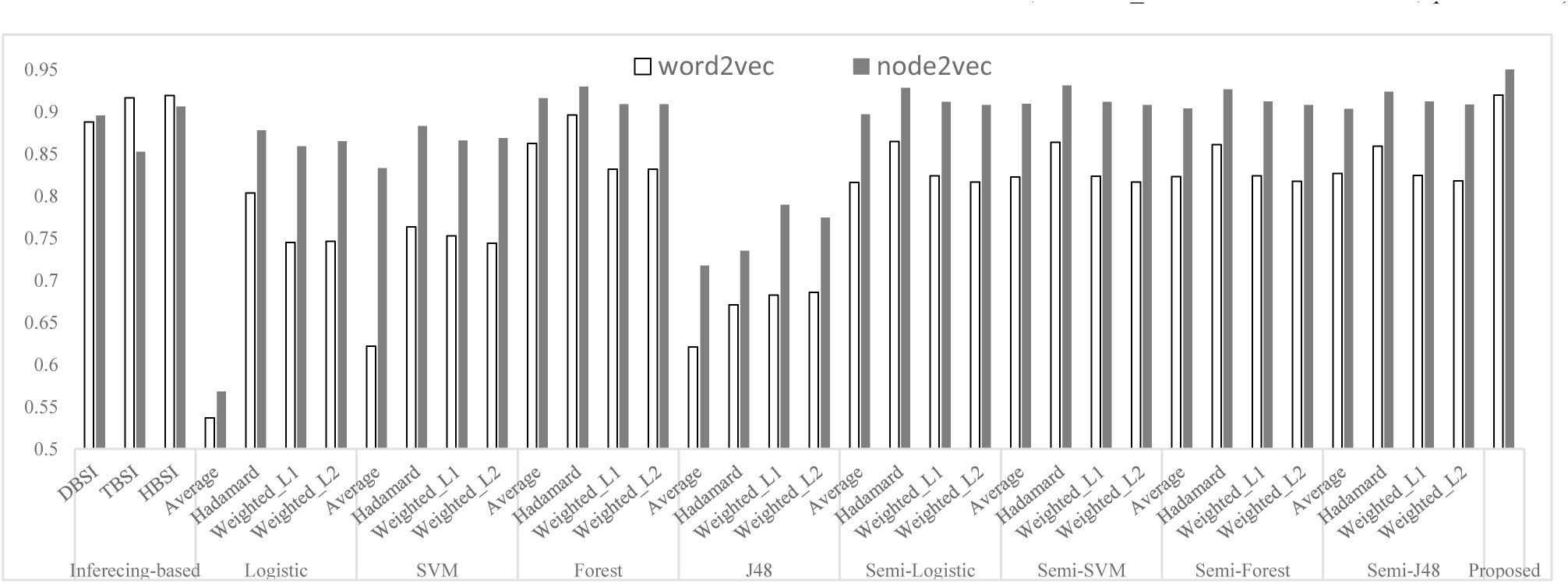
Comparison of average AUC scores. The hyper-parameter of DeepWalk was determined by a grid search over the parameter ranges specified in (Perozzi, et al., 2014) (number of walks *γ* = {10, 40}, learning rate *α* = {0.01, 0.05, 0.09}, dimension *d* = {64,128, 256}, window size *w* = {5, 10}, walk length *t* = {40, 80}). The hyper-parameter of Node2vec was determined by a grid search over the parameter ranges specified in (Perozzi, et al., 2014) (number of walks *γ* = {10, 40, Return *p* = {0.5, 1.0, 2.0}, in-out *q* = {0.5, 1.0, 2.0}, dimension *d* = {64, 128, 256}, window size *w* = {5, 10}, walk length *t* = {40, 80}). For the classification methods, the following settings are used as, L2 regularization for logistic regression, type C-SVC and kernel RBF for SVM, 500 trees for Random Forest, confidence factor 0.25 for J48.

Figure 1 also illustrates two phenomena: 1) the Node2vec-based methods show better results than Deepwalk-based (mean_Deepwalk=79.8%, variance_Deepwalk=0.8%, mean_Node2vec =87.5%, variance_Node2vec =0.6%, p=8.18 E-05); 2) the semi-classification-based methods outperform classification-based ones (mean_classification=79.2%, variance_classification=0.9%, mean_semi-classification =87.2%, variance_ semi-classification =0.2%, p=4.22 E-05).

### 3.2 Binary operator and classification methods for prediction

To understand the effect of the two important components for the association prediction model in the proposed method, we took all the possible combinations of the methods for the two components into consideration and demonstrate the results in Figure 2.

For the binary operator, Average performed the best (AUC ROC= 95.3%) and Hadamard performed the second (AUC ROC= 94.5%). Average performed the best for most of the cases except for the word2vec with semi-classification. Weighted-L1, and Weighted-L2 show the similar results for all the tests.

For the classification methods, SVM performed the best for Node2vec (AUC ROC= 95.3%) and Radom Forest performed the second (AUC ROC= 94.9%). For the semi-classification, only the vectorization methods are important.

In addition, both Figure 2 (a) and (b) illustrate classification-based solution outperformed the semi-classification-based solution (mean_classification=91.8%, variance_classification=0.06%, mean_semi-classification=89.8%, variance_ semi-classification =0.2%, p=0.013).

**Figure 2.**
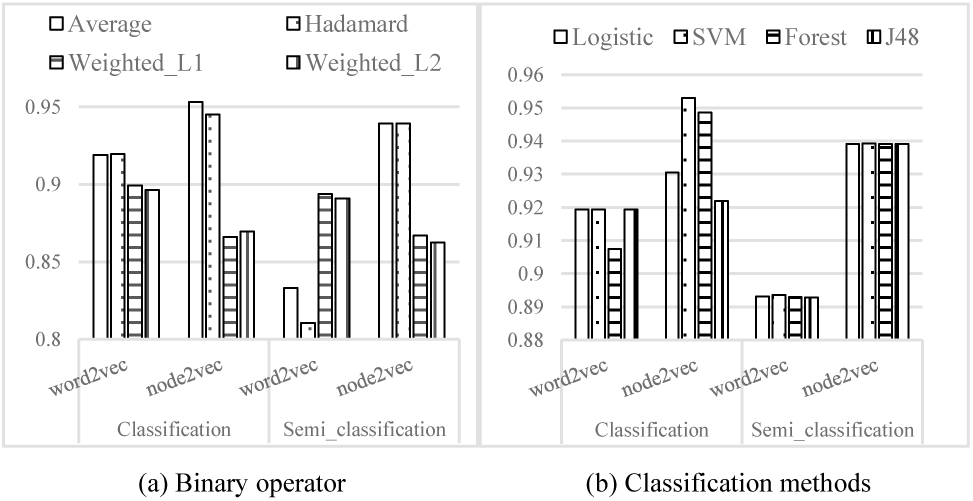
Comparison of methods for the two components in the proposed method.

### 3.3 Influence of negative samples

There are two kinds of associations, existent and nonexistent association, labeled as “positive” and “negative” and used in both training and testing. To distinguish those associations in all the data space, we classify the nodes into three kinds (See Supplementary figure 2), 1) Test Node Space (TNS), 2) Connected Node Space (CNS), 3) All Node Space (ANS). TNS is the space that contains the drug and target nodes used for testing. CNS is the space that contains the drug and target nodes having drug-target associations. ANS is the space that contains all the drug and target nodes. ANS includes CNS, and CNS includes TNS. Based on the three kinds of nodes defined, we can categorize all the associations into 7 types:

1. Test Node Associations (TNA), which are the associations that can be generated from a pair of drug and target from TNS.
2. Connected Node Associations (CNA), which are the associations that can be generated from a pair of drug and target from CNS excluding TNA.
3. All Node Associations (ANA), which are the associations that can be generated from a pair of drug and target from ANS excluding TNA and CNA.
4. TNA+CNA, which are the associations that can be generated from a pair of drug and target from CNS.
5. TNA+ANA, which are the associations that can be generated from a pair of drug and target from ANS excluding CNA.
6. CNA+ANA, which are associations that can be generated from a pair of drug and target from ANS excluding ANA.
7. TNA+CNA+ANA, which are associations that can be generated from a pair of drug and target from ANS.

To further understand and evaluate these associations affecting the predictions, we randomly selected negative associations from the 7 types for training and testing. We show the results of the 49 combinations in Figure 3.

The AUC ROC varies with different negative samples for training and testing, and the best AUC ROC can be achieved with ANA for training and testing, which almost improves 1.5% with TNA+ANA in Section 3.2. Compared to the existent associations (i.e., positive samples) in TNA for training, ANA provides the most distinguishable negative samples, which gives the best classification results. Figure 3 also gives the best training set for each kind of predictions test scenario, which the best training negatives lay in the diagonal of the table

**Figure 3.**
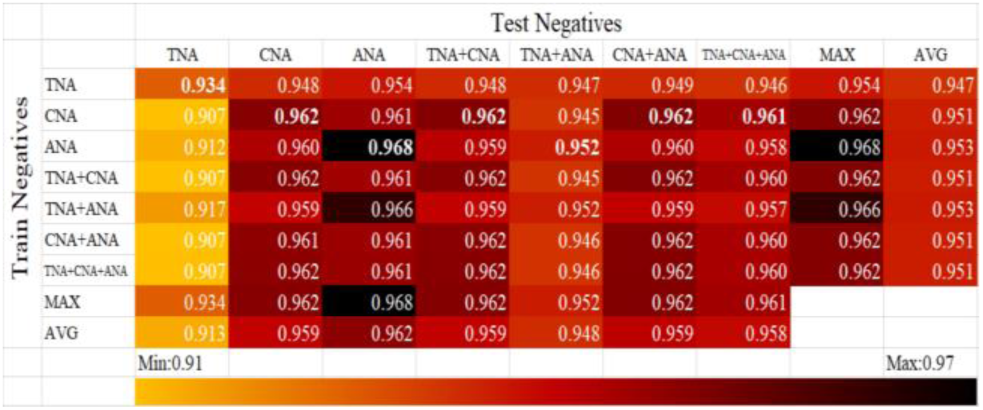
AUC scores of the prediction results of the proposed method with different combinations of training and testing negative samples.

**Figure 4.**
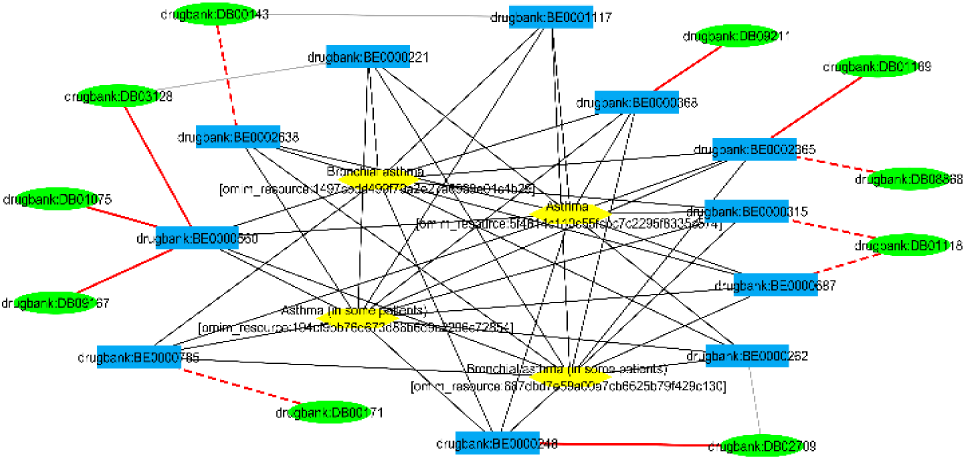
The Prediction for Asthma. Green ellipse and blue rectangle nodes are existent drugs and targets. Yellow ellipse nodes are disease entities for Asthma. The black solid lines are existent associations. The red solid lines are successfully predicted associations and the red dotted lines are failed one. The network file can be found in supplementary network 16.

**Figure 5.**
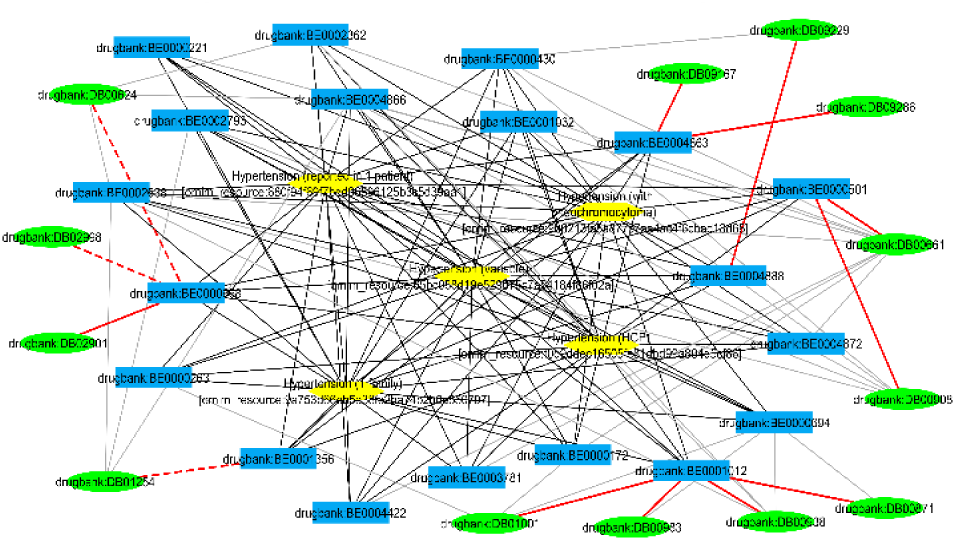
The Prediction for Hypertension. The network file can be found in supplementary network 19.

**Figure 6.**
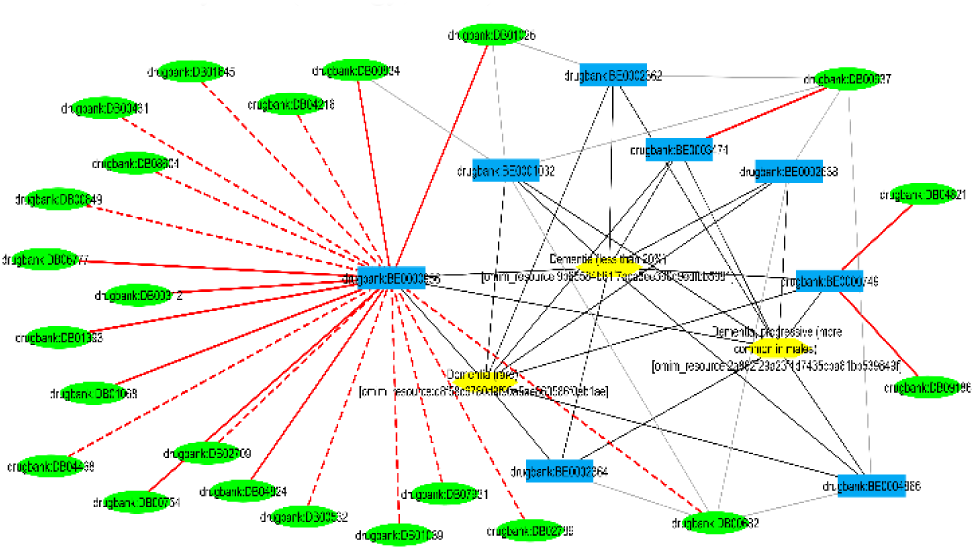
The Prediction for Dementia. The network file can be found in supplementary network 20.

## 4 Disease specific drug-target association prediction

With the parameters setting found in our best performed model in the previous experiment (*γ* = 10, *p* = 2.0, *q* = 1.0, *d* = 64, *w* = 5, *t* = 40), We trained our model with LMN, and try to find the novel drug-target associations for 20 diseases. For the positive associations, we searched DrugBank version 5.1.0 and validate them based on the drug-target 5.1.0. For the negative associations, we randomly selected 1000 times of positives in each of 7 types of training data introduced in Section 3.3. As Table 2 shows, there are totally 75 novel associations and 450,186 randomly selected negative samples (See Supplementary figure 3 for the screenshot of the predicted associations in the network and Supplementary network 1 for the data of the predictions).

**Table 2.** Disease specific drug-target prediction

| ID | Disease | # Positives | # Negatives | AUC ROC |
| --- | --- | --- | --- | --- |
| 1 | Adenocarcinoma | 1 | 6000 | 100.00% |
| 2 | Anemia | 1 | 6000 | 100.00% |
| 3 | Apert syndrome | 1 | 6000 | 100.00% |
| 4 | Autism | 1 | 6000 | 100.00% |
| 5 | Cirrhosis | 1 | 6000 | 100.00% |
| 6 | Galactosemia | 1 | 6000 | 100.00% |
| 7 | Insomnia | 1 | 6000 | 100.00% |
| 8 | Lung Cancer | 1 | 6000 | 100.00% |
| 9 | Malaria | 1 | 6000 | 100.00% |
| 10 | Obesity | 1 | 6000 | 100.00% |
| 11 | Obsessive Compulsive Disorder | 4 | 24000 | 100.00% |
| 12 | Spinal Muscular Atrophy | 1 | 6000 | 100.00% |
| 13 | Stroke | 4 | 24002 | 97.82% |
| 14 | Bipolar Disorder | 3 | 18003 | 96.99% |
| 15 | Asthma | 11 | 66069 | 96.03% |
| 16 | Adrenal Hyperplasia, Congenital | 2 | 12000 | 95.55% |
| 17 | Adrenal Hyperplasia, Congenital,<br>Due to 21-hydroxylase Deficiency | 2 | 12000 | 95.55% |
| 18 | Macular Degeneration | 1 | 6000 | 91.22% |
| 19 | Hypertension | 13 | 78065 | 90.93% |
| 20 | Dementia | 24 | 144047 | 80.36% |
| Total |  | 75 | 450186 | 97.22% (avg) |

The best AUC ROC scores are made for “Adenocarcinoma”, “Anemia”, “Apert syndrome”, “Autism”, “Cirrhosis”, “Galactosemia”, “Insomnia”, “Lung Cancer”, “Malaria”, “Obesity”, “Obsessive Compulsive Disorder”, and “Spinal Muscular Atrophy” (AUC ROC=100%). Other diseases received the AUC ROC scores ranged from 97.8% to 90.9%. The worst score is made for “Dementia” (AUC ROC=80.36%).

In addition, we received top K predictions from the proposed method and calculated the percentage of positives predicted (i.e., Recall). 50 associations are predicted over 75 positives (average 81.2% recall for each disease) with top 0.1% predictions (See Supplementary figure 4). The summary and detail information of top K arranged from 0.01% to 1% for each disease can be found in the supplementary Table 1 and 2.

The predictions for hypertension, dementia, and asthma across 20 diseases are the most prevalent results (All the disease specific predictions can be found in the supplementary networks 2-20 except for the disease “Macular Degeneration”, where the prediction fails in top 0.1%). These three diseases computed the greatest number of positives and were identified the meaningful targets that are more favorable for prediction in our model.

Asthma is a common chronic condition in which the airways become inflamed and narrowed. Patients living with this chronic diseases will often have trouble breathing and may experience asthma attack, where symptoms can worsen from physical or environmental triggers. Asthma is often treated with a combination of controller and rescue inhalers, along with other drugs taken orally for more severe cases. As researchers have learned more about the condition, treatment options for practitioners continue to evolve. Studies in asthma have focused beyond the small airways and looked upon how to specifically target inflammation for asthma management (Holgate, et al., 1996).

In our model, the target muscarinic acetylcholine receptor M2 produced the most successful predictions in the asthma case. In this case, muscarinic acetylcholine receptor M2 possessed the second-most drug relations. The target, Cytochrome P450 3A4, had the most drug relations. However, according to DrugBank, most of the existing drugs that bind to muscarinic acetylcholine receptor M2 function as antagonists, whereas about half of the existing drugs that bind to Cytochrome P450 3A4 function as substrates. We have found that the predictions for asthma that were based on the data for muscarinic acetylcholine receptor M2, which has mainly antagonist drugs, have performed better compared to predictions for other targets.

Muscarinic acetylcholine receptor M2 have contributed to three out of six successful predictions with through our model. The muscarinic acetylcholine receptors are strongly associated to the physiological effects of the airways because they intermediate the effects of parasympathetic nerves (Coulson and Fryer, 2003). The autonomic controls of airway are mainly controlled by the release of acetylcholine from the parasympathetic nerves. Particularly, M2 muscarinic receptors are mainly located on airway smooth muscles (Haddad, et al., 1994). A recent study has shown that the loss of M2 muscarinic receptors will increase the release of acetylcholine from the parasympathetic nerves which will lead to increase airway hyperreactivity (Coulson and Fryer, 2003). When there is an increased release of acetylcholine, muscarinic acetylcholine receptors are stimulated to contract smooth airway muscles. Activated M2 receptor will couple to G-protein Gi, in which it will produce contraction of the airway smooth muscle and hinder relaxation (Hirshman, et al., 1999). The effect of inconsistent and severe airway smooth muscle contractions may result in asthma attacks for asthma patients.

The second type of disease with the most positives is hypertension. Hypertension is medically considered as a condition with a sustained increase of blood pressure (BP) beyond 140 mmHg (systolic) over 90 mmHg (diastolic). It is the major cause of cardiovascular and renal diseases and strokes (Panchanatham and Shah, 2014). In the last decade, antihypertensive medications prescribed in various oral tablet combinations along with diet and exercise helped to control blood pressure. Antihypertensive medications can be separated into multiple types based on their mechanism of action, which include diuretics, beta-blockers, alpha blockers, angiotensin-converting enzyme inhibitor (ACE), and angiotensin receptor blockers (ARB). The management of hypertension often requires more than one or two drugs to efficiently control blood pressure over a lifetime (Panchanatham and Shah, 2014).

Using the data retrieved from DrugBank, there is a total of 116 drug relations for alpha adrenergic receptors and 22 drug relations for beta-3 adrenergic receptors. The number of drug relations from both alpha-adrenergic receptors and beta-3 adrenergic receptors is the most compared to the number of drug relations from other targets. Additionally, we have also found that existing drugs that bind to these targets mainly serves as either antagonist or agonist. Thus, with the greatest number of drug relations, which were mainly associated with antagonist and agonists drugs, beta-3 adrenergic receptors, alpha-1, alpha-1A, and alpha-adrenergic receptors predicted the greatest number of positives in hypertension through our model.

Four out of five targets possessing successful predictions are alpha-1, alpha-1A, alpha, and beta-3 adrenergic receptors. Both alpha- and beta-3 adrenergic receptors are associated to muscles contractions. Drugs that are predicted mainly by beta-3 adrenergic receptors can be classified as beta-blockers. Beta blockers’ mechanism of actions includes blocking the hormone epinephrine, or adrenaline, from taking effect, which in turn reduce the rate and force the heart uses to pump blood, therefore, lowering the blood pressure. In addition, drugs that are mainly predicted by alpha-adrenergic receptors can be classified as alpha blockers. Alpha blockers keep the hormone norepinephrine through tightening the muscles in the walls of veins and small arteries. Thus, they can lower blood pressure as they keep blood vessels open and allow blood to flow more efficiently. Our model was able to successfully predict the drugs related to these receptors by using their drug relations as the adrenergic receptor family that mediates the sympathetic nervous system and regulate the cardiovascular system (Kanagy, 2005).

Finally, the third type of disease with the most positives is dementia. Aging and elderly adults commonly suffer from the decline in their mental ability. Dementia, a severe decline in cognitive function that tremendously impacts a person’s daily life, is a common cause of cognitive impairment in elderly (Kivipelto, et al., 2018). Dementia is a neurocognitive disorder in which the cognitive functions such as learning and memory, languages, social cognition are severely impaired. The potential of one getting dementia increases exponentially from the age 65 to 85 years (Jellinger and Attems, 2010). According to the causes, dementia can be classified into various types such as vascular dementia, frontotemporal dementia, and Alzheimer’s disease (Kaur, et al., 2018).

Similar to the results from hypertension, the target, nuclear receptor subfamily 1 group I member 2, or pregnane X receptor (PXR), which has the most successfully predictions in our model, has the second-most drug relations. Additionally, according to DrugBank, the functions for most of the existing drugs that bind to PXR are distributed across the categories of antagonist, agonist, unknown, activator, binder, partial agonist, suppressor, and inducer. Thus, with the greatest number of drug relations, PXR predicted the greatest number of positives in hypertension through our model.

PXR was able to predict seven out of ten drugs within our model. The numbers of predictions that can be made from this gene alone indicate its potential to predict new drugs for dementia through our method. PXR is a member of nuclear receptor and is generally expressed in liver and intestine. It is usually associated to the regulation of enzymes and transporters that are responsible for metabolism and removals of xenobiotics and endobiotic (Ma, et al., 2008). In addition, recent studies have shown that PXR may serve as a fairly attractive target in treating for dementia due to its role in cognitive deficits (Kaur, et al., 2018).

## 5 Discussion and Conclusion

Despite the reliability of the proposed method proven in this study, there were a couple noted issues we will disclose in order to be enhanced in the future.

Firstly, the topology of the network is used to obtain the vector representations (i.e., features) of the vertices. The network is treated flat and all the edges are considered equal. However, the biomedical linked datasets and real world heterogeneous networks contain diverse properties to describe the knowledge and data, which is difficult modelled in a single flat network (i.e., one dimensional network) interpretable. Using one vertex to model same entities for diverse context will cause the relations computational undistinguishable. The solution used in this study is to create separate nodes for the same concept, and copy all the linkages to the new twin nodes. However, the solution used can increase computation. A more straightforward solution is to apply multiple-dimension weighted graph. However, to the best knowledge for the authors, there lacks the solid embedding solution for multiple-dimension weighted graph nor it is not feasible to weight the properties. A simpler solution is to embed multiple-dimensional graph (Ma, et al., 2018; Ma, et al., 2018). Instead of integrating heterogeneous networks to obtain a network for feature learning, vector fusion methods can be applied to formalize a node vector based on the vectors learned from each heterogeneous network (Luo, et al., 2017).

Secondly, the feature learning and classification are two separate steps in classification-based drug-target prediction models (Grover and Leskovec, 2016; Perozzi, et al., 2014; Tang, et al., 2015). Feature learning plays important role to the quality of the final classification results, but not vice versa. Therefore, suitable feature learning method and parameter tuning may dramatically affect the final prediction (Zong, et al., 2017), which brings more potential risks to compromise the results. One potential solution is to train the feature learning and classification models together in a prediction model and allows feedback mechanism to boost the parameter learning for feature learning (Feng, et al., 2018), which can be built based on the idea of Deep Neural Networks.

Lastly, one of the main objective for the computational drug-target prediction methods is to narrow down the screening candidate for in vitro experiments, such as drug-repurposing. Therefore, in practical applications, with a list of given drugs or targets as the input, the whole targets or drugs in the database should be considered as the potential candidate in prediction. The big imbalanced test data in pairwise prediction will cause the problems of feasibility in computation and bias in result. Although we have discussed the strategy of selecting the pairs in training, we did not give the efficient strategy to deal with the huge number of candidate pairs in testing when those pairs are coming from different groups and imbalanced. One of the solutions is to add a search function (i.e., ranking function) to pre-select the candidate pair for testing. Different with the negative selection method we applied to deal with given network topology (Liu, et al., 2002), the search method for test candidates should give dynamic pairs based on the inputs.

In conclusion, we provided a network-based solution that predicts novel drug-target associations based on the integrated results from an inference-based model and a classification model. The features used in the two models are generated based on a heterogeneous network including #20,119 entities, # 194,296 associations. With the 10-fold cross validation experiments and a disease specific prediction task based on 20 diseases, we showed that the proposed method can be used as a reliable method to identify novel drug-target associations.

## Acknowledgments

The authors appreciate the valuable opinion from Dr. Rui Chen. The authors also thank Dr. Sejin Nam to provide the computational support.

## Footnotes

1 DrugBank, Human Disease Network, Sider, KEGG, PharmGKB, Pubchem, OMIM, Uniprot, HGNC, UMLS, MESH, Bio2rdf

2 Drug, Target, Disease, Sider effect, Pathway, Haplotype, Variant location

3 1) Adenocarcinoma, 2) Anemia, 3) Apert syndrome, 4) Autism, 5) Cirrhosis, 6) Galactosemia, 7) Insomnia, 8) Lung Cancer, 9) Malaria, 10) Obesity, 11) Obsessive Compulsive Disorder, 12) Spinal Muscular Atrophy, 13) Stroke, 14) Bipolar Disorder, 15) Asthma, 16) Adrenal Hyperplasia, Congenital, 17) Adrenal Hyperplasia, Congenital, Due to 21-hydroxylase Deficiency, 18) Macular Degeneration, 19) Hypertension, 20) Dementia

